# Slow and steady wins the race: encounters of myosin-5 and myosin-6 on shared actin filaments

**DOI:** 10.1101/087940

**Authors:** Alicja Santos, Joanna Kalita, Ronald S. Rock

**Author notes:** Corresponding Author: Ronald S. Rock, GCIS W240, 929 E. 57th Street, Chicago, IL 60637.

## Abstract

In the cellular environment multiple myosins use the same filamentous actin (F-actin) tracks, yet little is known about how this track sharing is achieved and maintained. To assess the influence that different myosin classes have on each other, we developed an assay that combines two dynamic elements: elongating actin filaments with identified barbed and pointed ends, and myosins moving along these filaments. We studied two different myosins, myosin-5 and myosin-6. These myosins have distinct functions in the cell and are known to travel in opposite directions along actin filaments. Myosin-5 walks towards the barbed end of F-actin and generally into dynamically rearranging actin at the cell periphery. Myosin-6 is a pointed-end directed myosin that generally walks towards the cell center. We successfully reconstituted simultaneous bidirectional motility of myosin-5 and myosin-6 on single polymerizing filaments of actin. We report and provide statistical analysis of encounters between myosin-5 and myosin-6 walking along the single filaments. When myosin-5 and myosin-6 collide, myosin-5 detaches more frequently than myosin-6. The experimental observations are consistent with a stochastic stepping model based upon known myosin kinetics, which suggests that faster motors are more likely to detach.

---

Myosin motor classes co-exist in the cell and use the same F-actin tracks to transport material and to apply forces to the cytoskeleton. It is currently unclear how these myosins react when they encounter other myosins or obstacles, even though such encounters can be common in the cell. The F-actin networks have different composition in subcellular regions, creating distinct local environments [1]. In some areas of the cell, where F-actin tracks are limiting, myosins could crowd onto single tracks. Determining the accessible myosin binding sites on actin and the outcome when myosins compete for the same site reveals important aspects of self-organization in the eukaryotic cytoskeleton.

In our previous work we used assembling F-actin to test how actin tracks modulate myosin motility. We discovered that myosin-5 and myosin-6 detect the age of actin tracks via the actin nucleotide state, and adjust their processivity accordingly [2]. Myosin-5 has longer runs on young filaments, corresponding to leading edge in the cell. Myosin-6 has longer runs on old filaments, as found in the cell interior away from sites of filament nucleation. Here, we consider another aspect of myosin-5 and myosin-6 motility that has been omitted in the *in vitro* assays. Because myosin-5 and myosin-6 travel in opposite directions, they should encounter each other frequently and eventually compete for the same binding site along actin.

In this work we seek to answer two main questions about myosin-5 and myosin-6 motility. The first is whether a single F-actin track supports the simultaneous traffic of myosins traveling in opposite directions. The second is how such opposing motility is resolved to ensure effective myosin-5 and myosin-6 based transport.

The main technical challenge in reconstituting the motility of oppositely directed myosins is establishing the polarity of the F-actin. Several groups have devised approaches to distinguish the barbed and pointed ends of actin filaments in a motility assay. The main approach is to use an anchoring particle on one end of F-actin, followed by solution flow to orient filaments in the image plane of the microscope [3-5]. Only one of these studies examined motility of myosin-5 and myosin-6 on the same filament [5]. This study by Yuan *et al*. focused on the shear flow methodology with anchored gelsolin used to orient the filaments. These relatively elaborate preparation challenges resulted in a low number of myosin-5 / myosin-6 encounters (n=26). Moreover, Yuan *et al*. used quantum dots to label their myosin motors. Here we are interested in how the myosin motor domains compete for the same binding-sites along actin, and how this competition is resolved. To examine these features of motor-domain competition, it is important to avoid bulky fluorophores like quantum dots (QDots), which may present steric interference and affect the processivity of the myosins. On the other hand, quantum dots would mimic the bulk of a cellular cargo, so each approach has its own merits.

Here, we monitor myosins on polymerizing F-actin as in our prior work [2]. The rapidly elongating end of actin is the barbed end. Myosins that travel toward the elongating end are myosin-5s [6], and those that travel away from the elongating end are myosin-6s [7]. This system does not require flow to align the filaments, nor does it need an anchor or a fluorescent marker to cap the barbed end. We use the same fluorophore on each myosin: a small, Cy5-labeled affinity clamp [8]. The affinity clamp is comparable in size to a GFP label and much smaller than a QDot, reducing steric interference [5, 8, 9]. This labeling approach also simplifies the imaging conditions and ensures equivalent labeling of each myosin.

We successfully reconstituted simultaneous motility of myosin-5 and myosin-6 on single polymerizing filaments (Figure 1a-c). By observing the elongation of actin from the start of the experiment, we can ensure that myosin-5 and myosin-6 travel along the same single F-actin. We easily identify barbed and pointed ends of each filament and the myosin identity for each myosin run (Supplementary Movie 1-2). Because myosin-5 and myosin-6 move at different speeds, we can use the speed to verify the myosin class assignment. The assignments agreed in every case. In Figure 1c we show representative kymographs of bidirectional motility. The diagonal front showing F-actin elongation is marked in red, and diagonal myosin paths are shown in green. Myosin-5 moves faster than myosin-6, resulting in a diagonal line that appears thinner despite carrying the same fluorophore. To optimize the number of myosin encounters, we increased the concentration of each myosin from 1 nM to 5 nM. A comparison of myosin density between the 1 nM and 5 nM in the presence of TMR phalloidin conditions is shown in the kymographs (Figure 1c). This 5-fold increase in concentration resulted in a 25-fold increase in the encounter frequency, from n = 21 for 1 nM myosin to n = 500 for 5 nM myosin, as expected for a first order process that is first order in each myosin.

**Figure 1.**
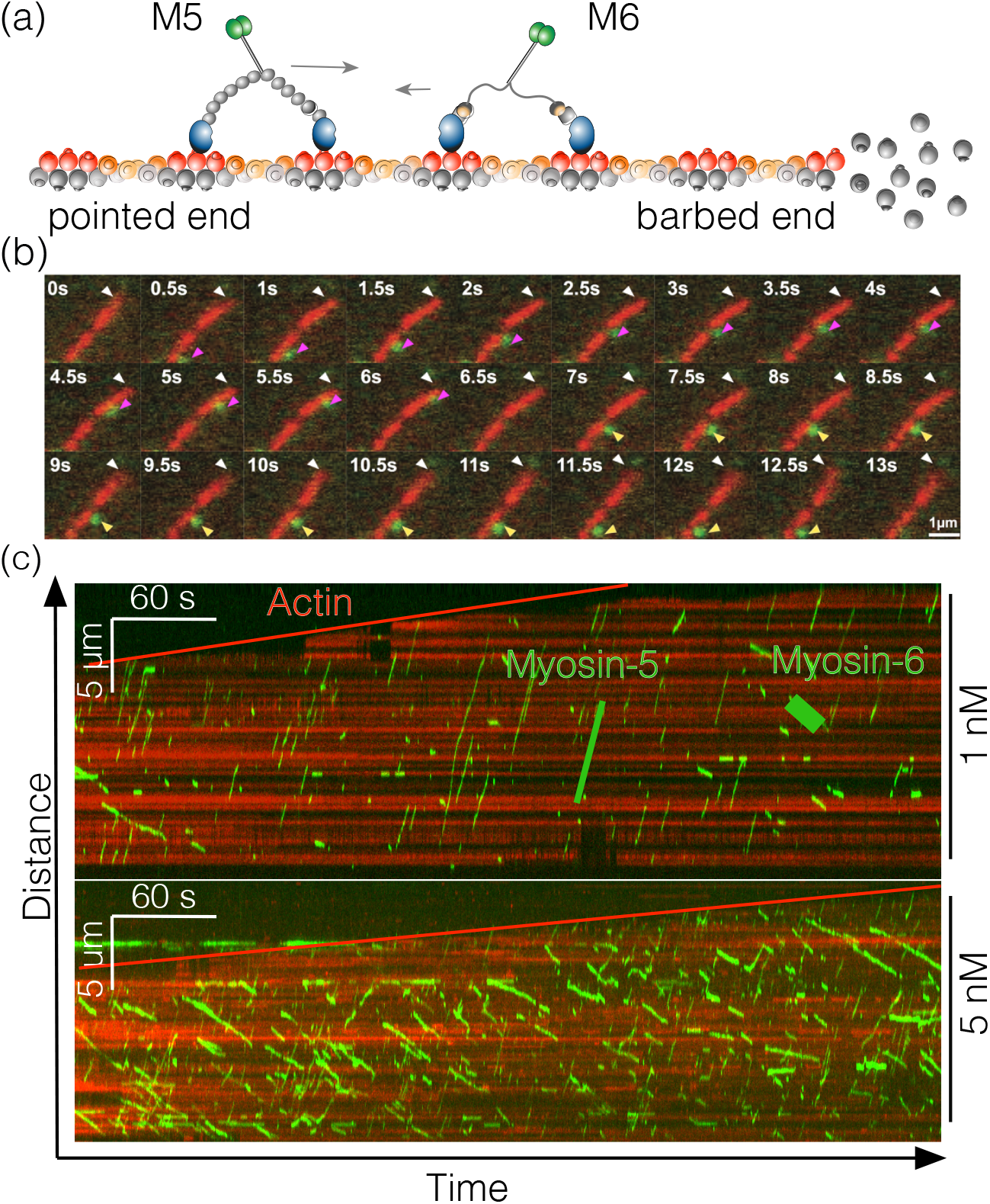
Myosin-5 and myosin-6 walk in opposite directions on the same filament *in vitro*. (a) Schematic of the experimental setup. Fluorescently labeled myosin-5 and myosin-6 walk along an assembling actin filament. (b) Time-lapse fluorescence micrographs of a single myosin-5 and a single myosin-6 (both green) moving along a single growing actin filament (red). White arrowheads mark the barbed end of the filament, growing to the upper-right corner. Magenta arrowheads mark a single myosin-5 traveling towards the growing end, and yellow arrowheads mark a single myosin-6 traveling away from the growing end. (c) Representative kymographs showing processive motility of myosin-5 and myosin-6 on assembling filaments. Although myosin-5 and myosin-6 are labeled with the same fluorophore, they can be distinguished by the direction of their runs (indicated in the top kymograph). Myosin-5 and myosin-6 travel towards the barbed (top) and pointed ends (bottom), respectively. Kymographs show the relative motility of both myosins at equal motor concentrations of either 1 nM or 5 nM each.

We observed a total of 500 crossings at equimolar, 5 nM concentrations of myosin-5 and myosin-6 (Figure 2a). From the kymographs, we distinguish five kinds of outcomes (Figure 2b-f) and report the number of events observed for each of them (Figure 2g). The most common outcome is for both of the myosins to continue their runs, passing by each other on the opposite sides of a filament; such events account for 53% of all encounters. The second most common outcome is for at least one myosin to continue its run after the encounter; such events account for 42% of all encounters. Myosin-6 continues the run in 27% of total number of encounters, while myosin-5 continues in 15% of total number of encounters. Encounters that result in detachment of both myosins or stalls are rare, 5% and 0.4% respectively. We do not observe extended pauses at the site of an encounter, thought we might be limited by our exposure time of 500 ms. Myosin velocities and runlengths are unaffected and in agreement with literature values (Fig. S1-2, Table S1-2) [10].

**Figure 2.**
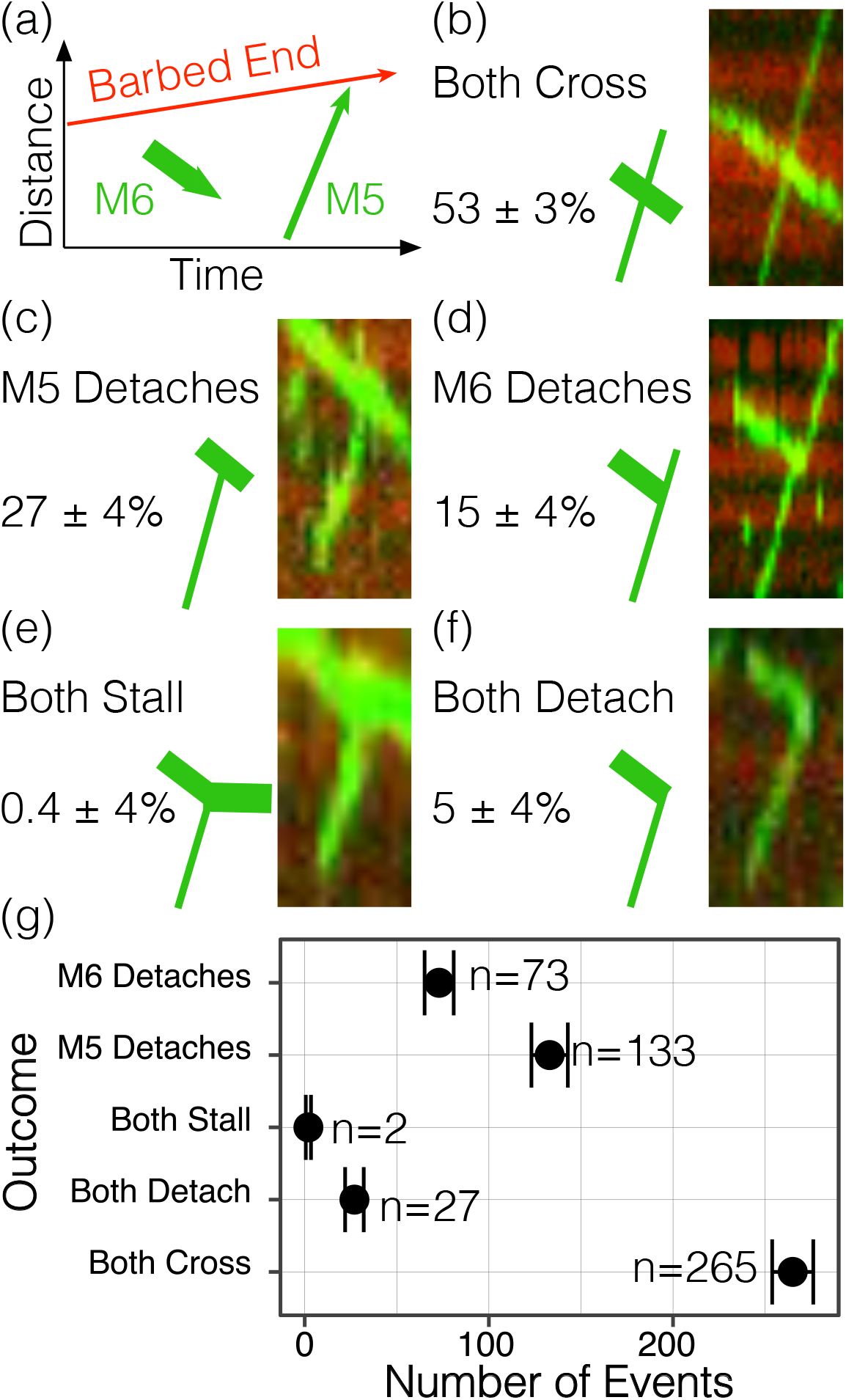
Collisions between myosin-5 and myosin-6 lead to five possible outcomes. (a) Scheme of kymograph analysis presented in (b-f). As in Figure 1, green diagonal lines from the lower left to the upper right are myosin-5, and lines from the upper right to the lower left are myosin-6. The red diagonal line represents the barbed end of growing actin filament. Directionality of actin growth and myosin motility are marked with arrows. Five possible collision outcomes are reported with percent values that they account for among total number of collisions (500, out of 2611 total myosin runs). (b) The majority of the encounters allow both motors to cross. (c-d) The second most common outcome is when one myosin continues while the other detaches at the site of the encounter. (e) Very few encounters result in both motors stalling, indicated by a horizontal green line after the encounter. (f) A few encounters end with both myosins detaching. (g) The frequency of collision outcomes. Among events where one of the myosins detaches and the other myosin continues the run, myosin-5 detaches first almost twice as often as myosin-6. Error bars show standard deviations estimated from multinomial counting statistics.

We can distinguish two broad behaviors of myosin-5 and myosin-6 walking toward each other on one filament. In half of the encounters, the myosins pass each other like “two ships that pass in the night”. The other half of encounters result in what we call “collisions,” where myosins interfere with each other at the encounter site. In a “head on” collision between myosin-5 and myosin-6, the two myosins compete for the same binding site on actin as they approach along the same path from opposite directions (Figure 3a-c). We use the term “collisions” here merely to indicate such a competition for binding sites, not to suggest a physical process involving an exchange of momentum.

**Figure 3.**
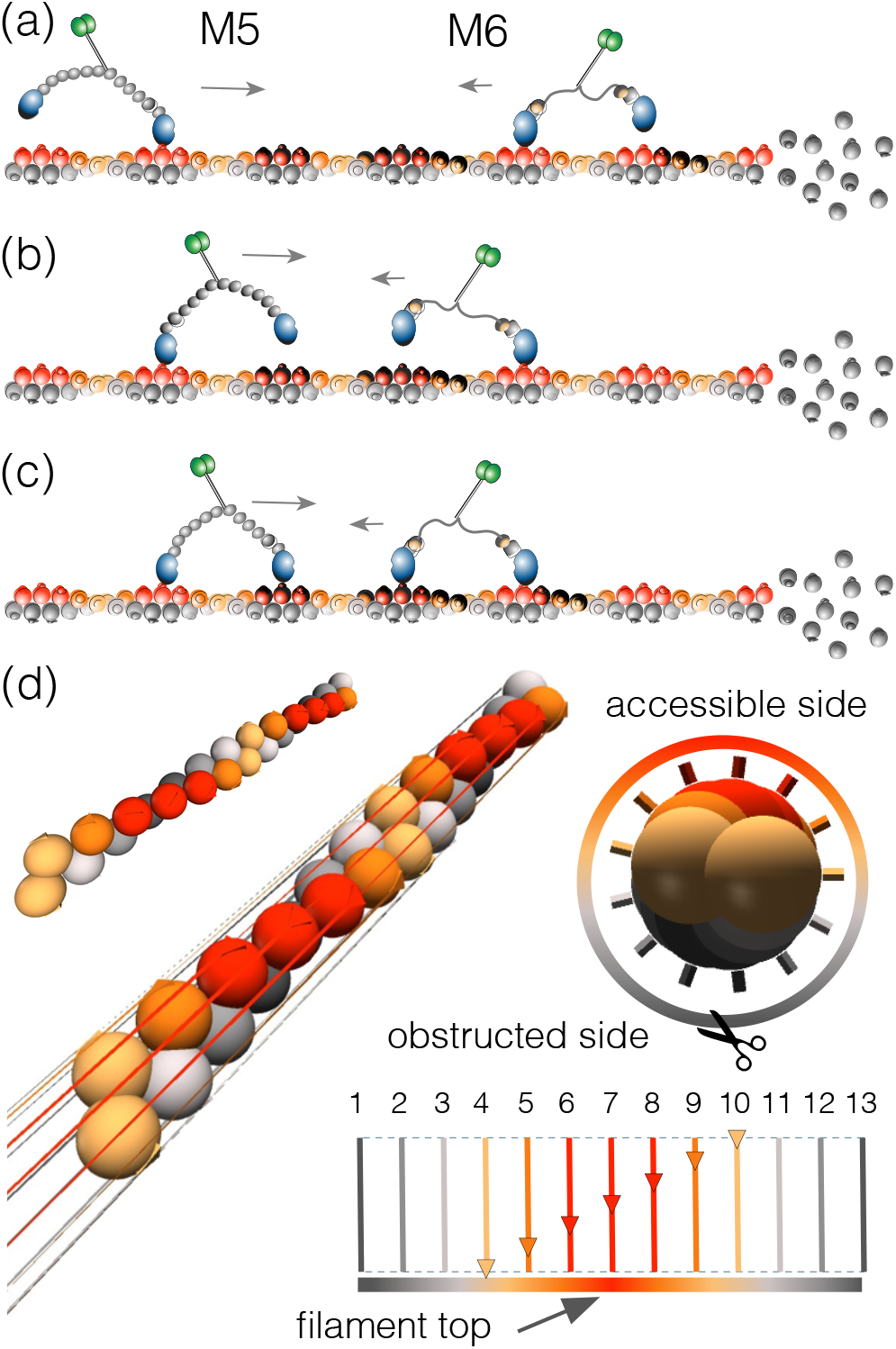
Spatial arrangement of myosin-5 and myosin-6 during bidirectional motility along a single filament. (a-c) Stages of myosin-5 and myosin-6 approaching a collision event on a growing actin filament, (a) Myosin-5 and myosin-6 are aligned along the top of the filament, within the plane of the page, and will eventually compete for the same actin-binding site If they continue along those paths without sidesteps. Accessible binding sites on actin are emphasized by darkened monomers. Myosin-5 uses one of three accessible actin monomers for forward steps, with a strong bias toward the central monomer of the three. Myosin-6 uses one of approximately six accessible monomers for the forward step and one of three accessible monomers for the backward step. (b) Myosin-5 and myosin-6 both complete their working strokes, positioning their free motor domains near their next target binding sites along actin. (c) Myosin-5 and myosin-6 rebind to the filament. With this alignment of the two myosins, one of the motors must detach in the next cycle unless a sidestep occurs and the lane is cleared. (d) Visualization of accessible tracks for myosin walking along the filament. There are thirteen lanes encircling the filament, shown as lines connecting consecutive monomers along each lane in the left panel. A slice looking down the long axis of the filament is shown in the right panel. At the bottom, the unfurled cylinder is shown with its thirteen lanes. The red lanes are easily accessible for myosin binding, orange and yellow are less exposed, and gray indicate actin lanes that are likely obstructed by the coverslip surface.

Passing events occur somewhat less frequently than expected considering the actin filament geometry. The actin filament forms a 13/6 helix, with 13 actin monomers per pseudohelical repeat. Thus, we expect that F-actin would present a maximum of 13 separate lanes for myosin traffic (Figure 3d). Filaments in the experiment are attached to the coverslip surface through streptavidin links. Even though the attachment sites are sparse, with actin biotinylation at 0.1%, we expect that half of the lanes are occluded due to proximity to the glass (Figure 3d). If seven F-actin lanes are accessible, we would predict that under 14% of runs would result in a collision and 86% of myosins would pass each other at the encounter site. Collisions might be more prevalent than expected under this model if each myosin occludes more than one lane, if myosin landing events are biased toward a smaller number of more exposed lanes (perhaps the ones farthest from the coverslip) [11], or through a combination of these two mechanisms.

Myosin-6 can take occasional (10%) backsteps and has a broad range of target binding sites along actin [12]. We expect that these backsteps could affect collision outcomes by up to 10%, depending on the detailed mechanism of backstepping. Myosin-5 has one preferred actin binding site when it takes a forward step, however it can bind one actin monomer before or after the preferred binding site with lower probability [13]. Stepping off the preferred binding site will cause the two myosins change lanes. However, sidesteps that avoid a collision are likely balanced by sidesteps that lead to a collision. Therefore we expect that sidesteps and backsteps have a negligible effect on the stepping outcome, and we omit an explicit treatment of either here.

Just under half of the cases are collisions, where the myosins appear to interact at the encounter site. Collisions that result in detachment of both myosins are rare (5%) and can be a result of myosins ending the run right after an encounter, despite presence or absence of the opponent. Based upon runlength estimates for these myosins, we predict that 5.5% of encounters would have both myosins detach within a 250 nm window, a reasonable limit on our ability to resolve passing events. Collisions that result in stall are extremely rare (0.4%) and could be an effect of the myosins entangling with each other. The bulk of collisions result in one of the myosins detaching at the encounter site. The most straightforward explanation is that one myosin occupies actin binding sites required by the other, in the head on collision scenario (Figure 3a-c). In 65% of these cases, myosin-6 continues the run while myosin-5 detaches. From our measured runlengths, we can determine that 15% of myosin-5 and 37% of myosin-6 will detach within our 250 nm resolution limit. We can correct our collision frequency by subtracting these false collisions, yielding an estimate that myosin-6 continues and myosin-5 detaches in 71% of collisions. We expect that this collision outcome is determined by a kinetic competition between each myosin for sites along actin.

To further understand the nature of this competition, we performed a stochastic simulation of the approach process of the two myosins. Briefly, we describe a set of lattice sites that mark myosin positions on three actin monomers in one lane along the filament. The myosin state is represented at the lattice site for its most advanced motor domain, where the free (even) sites are always between the bound (odd) sites (Figure 4a). We set transition rates between these lattice sites to experimentally determined myosin-5 and myosin-6 values for ADP release (myosin detachment), and Pi release (myosin reattachment) [14-17] (Table S1). To sample the transition times between lattice sites we use Gillespie’s direct method. The end of each simulation run occurs when one of the two myosins detaches. In 73% [14, 15] or 81% [16, 17] of our simulated collisions, myosin-5 detaches first. Both values compare favorably with our corrected experimental value of 71%.

**Figure 4.**
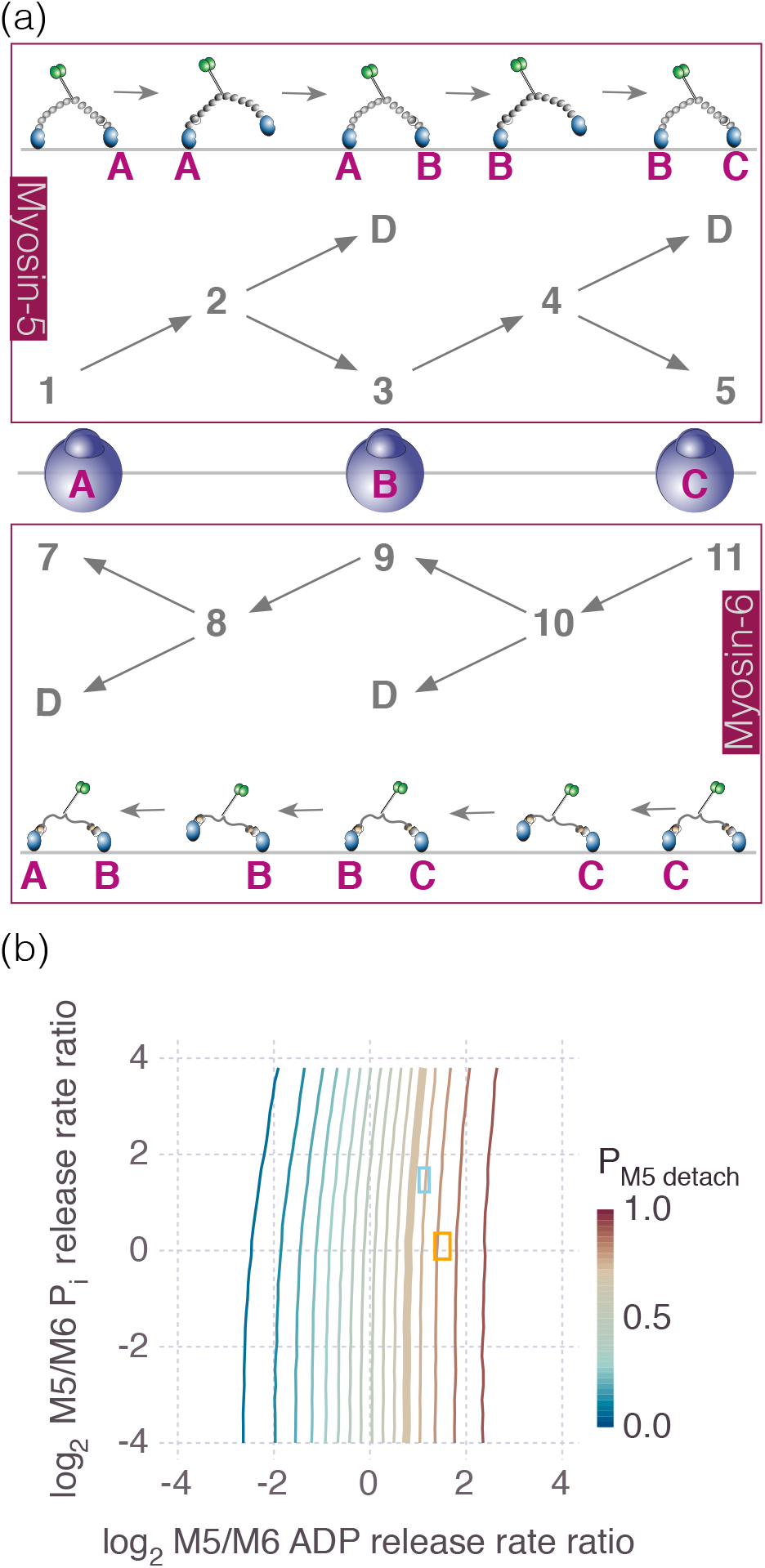
A stochastic simulation predicts that the faster myosin should detach more frequently when colliding with a slower myosin. (a) For simulation we consider three actin monomers along one path on filament as well as one myosin-5 molecule and one myosin-6 molecule that approach each other from opposite directions using these three monomers. Assigned numbers represent a set of lattice that mark myosin-5 and myosin-6 positions. The myosin state is represented at the lattice site for its most advanced motor domain, where the free (even) sites are always ahead of the bound (odd) sites. We use the experimental ADP release rates for the odd-to-even numbered transitions, and for the even-to-D transitions. Likewise, we use the experimental myosin phosphate release rates to approximate actin rebinding rates in the even-to-odd transitions. (b) A contour plot of the probability of myosin-5 detaching first at a collision, versus P_i_ release and ADP release rates ratios of myosin-5 to myosin-6. Rates are plotted as log_2_(k_m5_/k_m6_) values. Note that when both rates are equal (the point at the origin), each myosin is equally likely to detach as expected. Faster myosin-5 stepping rates (to the right) increase the myosin-5 detachment frequency, while faster myosin-5 attachment rates (to the top) decrease the myosin-5 detachment frequency. However, the effect is stronger for the former versus the latter. Experimental rates are indicated by boxes (blue box [14, 15]; orange box, [16, 17]). The boxes are sized to ± 1 SD in each dimension, reflecting the propagated experimental error on these parameters. The thick contour indicates our measured myosin-5 detachment probability. The proximity of the boxes indicates good agreement between our results and both sets of kinetic experiments.

We performed a sensitivity analysis to understand the relative importance of the kinetic parameters used in our simulation (Figure 4b). Higher myosin-5 stepping rates relative to myosin-6 (to the right on the ADP release axis) strongly increase the myosin-5 detachment probability. Likewise, higher myosin-5 attachment rates relative to myosin-6 (to the top on the Pi release axis) decrease the myosin-5 detachment probability, but to a lesser degree. For comparison, two sets of relative rate constants from the literature are indicated on Figure 4b. Although there is a range of literature values for myosin-6 attachment rates to actin [14-17], differences in this rate do not strongly affect our observed detachment probabilities.

We rationalize the collision outcomes as follows. The penultimate myosin state just prior to full detachment has only one motor domain attached to actin. The faster myosin is more likely to transition into that vulnerable state than the slower myosin in the same unit of time. Indeed, our simulations find that in the instant myosin-5 detaches, myosin-6 is more likely to have both motor domains attached to actin rather than only one (61% doubly-bound). Thus, the slower myosin is more likely to continue its run while the faster myosin is more likely to fall off. These effects are found on the horizontal axis of Figure 4b. Rarely, both myosins occupy singly-attached states with a gap of one lattice site between them. In this case, the faster myosin is more likely to outcompete the slower to continue its run. But as shown in Figure 4b, the detachment kinetics (ADP release) dominate the attachment kinetics (P_i_ release).

Myosin-5 and myosin-6 are both essential cellular transporters. When two way traffic occupies the same actin filament, these two myosin families would encounter each other and will occasionally occupy the same lane along a single actin filament. Efficient bidirectional transport would require one myosin to yield to the other in certain circumstances. An example of such a situation is found in sensory hair cells, where myosin-6 is thought to recycle and traffic membrane [18] as well as to anchor the plasma membrane at the base of the stereocilia [19-21]. Myosin-6 is highly concentrated at the narrow bases of the stereocilia with a limited number of actin filaments, while myosin-5 is absent from stereocilia and is enriched only in afferent nerve cells that innervate hair cells [22]. Yet in stereocilia, other myosins perform barbed-end directed transport, such as myosin-3 or myosin-15 [23, 24]. Microvilli in intestinal or renal epithelial cells are a related environment where two way traffic is maintained and both myosin-6 and myosin-5b are transporters [25-27].

Here we examined encounters of myosins with opposite directionality on the molecular level, and found that the faster myosin frequently loses in the competition for actin binding sites. Because myosin-6 is the only known myosin class that moves toward the pointed end of actin, slower walking might enable it to plow through the many opposing myosin classes that would otherwise force it off of actin. Simultaneous bidirectional motility gives us insights about how multiple myosin families share actin tracks. As such, our results help in understanding the regulation of cellular multi-myosin traffic. Because myosins recognize each other as obstacles on a filament, our results also address a general challenge of myosins encountering a stationary road block.

## Materials and Methods

Myosin-5 and myosin-6 constructs designed for these experiments are truncated versions of the molecules lacking globular cargo binding domains named Heavy MeroMyosins (HMM). In these constructs the coil-coiled region is fused to the GCN4-p1 leucine zipper to ensure dimerization [28]. The myosin constructs were expressed in Sf9 cells using baculovirus expression system as previously described and purified via Flag-tag affinity chromatography [29, 30]. The established Ctag / affinity clamp system was used for labeling each myosin monomer with single fluorophore [31]. For Ctag constructs, Cy5 fluorescent labeling was performed with separately expressed and labeled Clamp protein [8]. Clamp Cy5 labeling efficiency was 83%. Because each myosin has two clamp binding-sites, we estimate the percentage of unlabeled myosin as 3%. Actin was purified from chicken skeletal muscle as previously described [32] and was polymerized in the imaging chamber [2]. Three kinds of globular monomeric actin (G-actin) were used: ‘black’ unlabeled actin that contained the majority of the whole pool of protein, actin labeled with TMR for visualization of the polymer, and biotinylated actin to allow binding to the surface of a flow chamber coated with neutravidin. Actin labeling was performed with either TMR-maleimide or biotin-maleimide at Cys 374 as previously described [12].

For chamber preparation and single molecule imaging we used methods developed during actin age study [2]. Briefly, piranha cleaned and polyethylene glycol (PEG) coated coverslips and slides were assembled into flow cells by adhering to a microscope slide with double stick tape (chamber volume ≈10 μL). Flow chambers were coated with neutravidin (0.5 mg/mL) and then blocked with bovine serum albumin (1 mg/mL). We then perfused 1 μM total G-actin and 1 or 5 nM of fluorescently labeled myosin-5 and myosin-6. The assay buffer contained 10 mM imidazole (pH 7.0), 50 mM KCl, 1 mM MgCl^2^, 1 mM EGTA, 50 mM DTT, 0.2 mM ATP, 50 μM CaCl_2_, 15 mM glucose, 20 μg/mL catalase, 100 μg/mL glucose oxidase, and 0.5% 400 centipoise methylcellulose). The dynamic actin had final composition of 1 μM total concentration, with 0.1% biotinylated actin. Preliminary tests on reconstituting simultaneous motility of myosin-5 and myosin-6 were performed with use of TMR labeled actin at 5% (Supplementary movie 1). For collection large sets of data and evaluating collision outcomes TMR phalloidin was present, sustaining ADP•Pi state of the filament and ensuring bright labeling without any use of TMR actin (Supplementary movie 2).

Motor runs, filament ends and crossing events as well as collision outcomes were traced manually using ImageJ. Trajectory runlengths, velocities and landing events were extracted using custom scripts written in Julia [33].

To model head on collisions we applied Gillespie’s direct method, as implemented in Julia [34]. We consider three consecutive actin monomers on a single lane along the filament and describe a set of lattice sites that enumerate myosin states. Myosin-5 advances along sites 1→2→3→4→5, while myosin-6 advances along sites 11→10→9→8→7. Lattice site 6 is always empty, which maintains the convention that odd states are two-head-bound states, while even states are single-head-bound states. We also include two virtual lattice sites that record myosin-5 or myosin-6 detachment. Using separate sets of lattice sites for each myosin is a simple means to track each of them in the Gillespie algorithm. Note that site pairs (1, 7), (3, 9), and (5, 11) are equivalent. When an odd-numbered site is occupied by one myosin, the transition rate into the corresponding site for the other myosin is set to zero. Each odd site can transition to the next even site, while each even site can transition to the next odd site or to the detached virtual sites. We use experimentally determined rate values for ADP release to indicate detachment and Pi release to approximate rebinding from the experimentally determined values [14-17, 35]. Each simulation, including each of the 27x27 parameters examined in Figure 4b, recorded the outcomes for 100000 collisions. When we set equal rate constants for both myosins, the detachment probability of myosin-5 is 0.5 as expected (Figure 4b).

## Author Contributions

A. S. and R.S.R. designed the experiments. A.S. performed experiments. J.K. provided reagents and assistance. A.S. and R.S.R. analyzed the data and wrote the manuscript.

## Acknowledgments

The authors would like to thank helpful comments from Shohei Koide, Erin Adams, David Kovar, and Ed Munro. This work was supported by NIH R01 GM078450 and GM109863 (to R.S.R.), and American Heart Association Predoctoral Fellowship 16PRE26850006 (to A.S.).

